# More or less deadly? A mathematical model that predicts SARS-CoV-2 evolutionary direction

**DOI:** 10.1101/2022.03.10.483726

**Authors:** Zhaobin Xu, Qiangcheng Zeng

## Abstract

SARS-CoV-2 has caused tremendous deaths world wild. It is of great value to predict the evolutionary direction of SARS-CoV-2. In this paper, we proposed a novel mathematical model that could predict the evolutionary trend of SARS-CoV-2. We focus on the mutational effects on viral assembly capacity. A robust coarse-grained mathematical model is constructed to simulate the virus dynamics in the host body. Both virulence and transmissibility can be quantified in this model. The relationship between virulence and transmissibility can be simulated. A delicate equilibrium point that optimizing the transmissibility can be numerically obtained. Based on this model, we predict the virulence of SARS-CoV-2 might further decrease, accompanied by an enhancement of transmissibility. However, this trend is not continuous; its virulence will not disappear but remains at a relatively stable range. We can also explain the cross-species transmission phenomenon of certain RNA virus based on this model. A small-scale model which simulates the virus packing process is also proposed. It can be explained why a small number of mutations would lead to a significant divergence in clinical performance, both in the overall particle formation quantity and virulence. This research provides a mathematical attempt to elucidate the evolutionary driving force in RNA virus evolution.

## 1 Introduction

History indicates that no lethal virus pandemic last forever. For example, the 1918 flu, which claimed tens of millions of lives, was evolved into a significantly less deadly seasonal flu after 1920 [1-2]. Similar situation happened such as 2009 H1N1 virus gradually lose its virulence soon after the pandemic [3-4]. The pandemic caused by RNA virus is different from those caused by bacterium in a way that it normally engages a quick virulence declination after the outbreak. As far as scientist can tell, the bacterium that caused the black death never lost its virulence [5]. Now the question emerges: will SARS-CoV-2, the virus that causes COVID-19, follow a similar trajectory? So far as we observe, there is a significantly declination of SARS-CoV-2’s virulence. For instance, the virulence of omicron strain is less than the delta strain, and the virulence of delta strain is less than the original strain. However, lots of people concerns the emergence of a more virulence strain. It is useful when we refer to the history but history itself is also highly unreliable. Therefore, it is necessary to derive a scientific theory that can help predict the evolutionary direction of SARS-CoV-2.

The scientific exploration toward this question started in early 1980s, when two mathematical biologist Roy Anderson and Robert May, proposed that germs transmit best when hosts shed a lot of the pathogen [6-7]. virulence and transmissibility go hand in hand, until the germ gets so deadly it winds up killing its host too soon, and therefore can’t spread at all. This is known as the transmission-virulence trade-off [8-10]. The most familiar example is that of [the myxoma virus](https://www.pnas.org/content/114/35/9397), a pathogen introduced to Australia in 1950 to rid the country of rabbits. Initially, the virus killed more than 90 percent of Australian rabbits it infected. But over time, a tense truce developed: Rabbits evolved resistance, the myxoma germ declined in virulence, and both rabbits and germ remained in precarious balance for some time [11].

A second theory, developed by evolutionary epidemiologist Paul Ewald, which he calls the “theory of virulence,” suggests that, as a rule, the deadlier the germ, the less likely it is to spread [12]. The reason: If victims are quickly immobilized (think of Ebola, for example), then they can’t readily spread the infection. By this thinking, if a germ requires a mobile host to spread, its virulence will, of necessity, decline. Like the older conventional wisdom, the theory of virulence recognizes that many germs will evolve less virulence as they circulate and adapt to the human population. But Ewald’s theory also proposes that germs all have their own strategies to spread, and some of those strategies allow the germ to maintain high virulence and transmissibility.

Here we proposed a new mathematical theory that predicts the virus evolutionary direction. Instead of using logic language and examples, we developed a mathematical approach that predict the evolutionary trend of RNA virus. We focus the effect of mutation on viral assembly capacity and explain why the alternation of viral assembly capacity will significantly change its virulence and transmissibility.

The viral assembly capacity has been extensively studied [13-20]. However, little attention is paid to linking this assembly kinetics into its virulence evolution. In the result section 3.1, we first illustrate why and how this self-assembly capacity influence its replication and the formation of virus particles. In 3.2 we apply this model to predict the delicate equilibrium point between virulence and transmissibility, which is also the point that maximizes the transmissibility. In 3.3, we explain why a tiny number of mutations could lead to a significant difference in virulence and overall packed virus particles. This theory provides some possible explanations toward those interesting questions below

I:What is the exact relationship between virulence and transmissibility?

II: Do we need to worry about the emergence of a more deadly strain?

III: Will the virulence of SARS-CoV-2 finally fade away?

IV: Why the cross-species transmission always leads to a fatal outbreak in the early epidemic stage?

V:Why a few mutations can significantly alter the clinical behaviors of SARS-CoV-2?

## 2 Methods

### 2.1: Introduction of our coarse-grained model (model 1)

We used ordinary differential equations to describe the dynamics of the virus together with antibodies. 14 reactions are constructed:

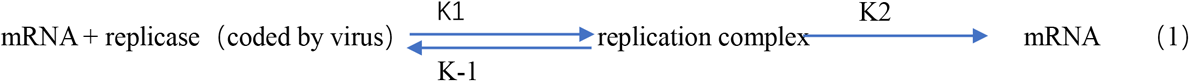

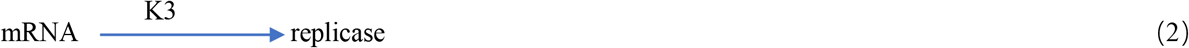

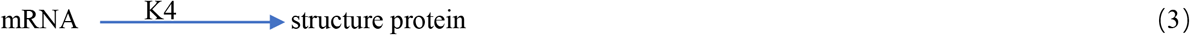

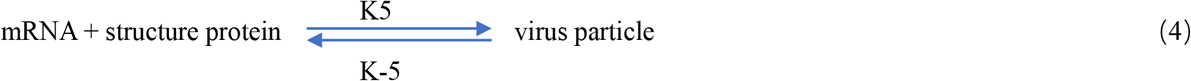

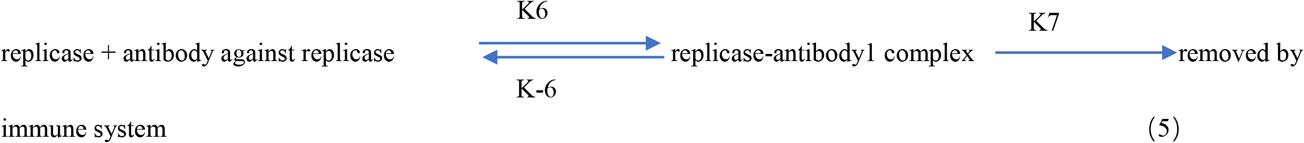

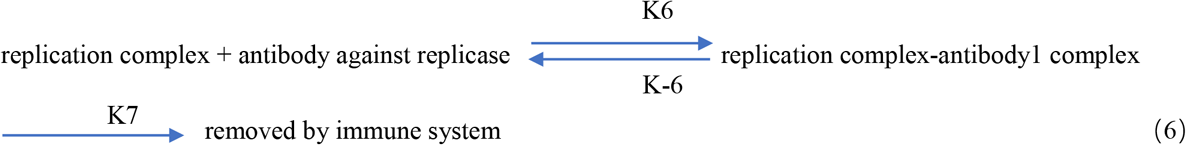

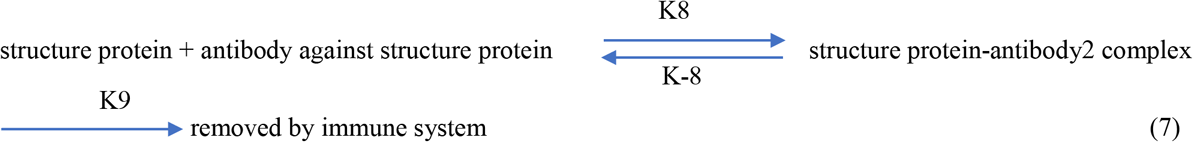

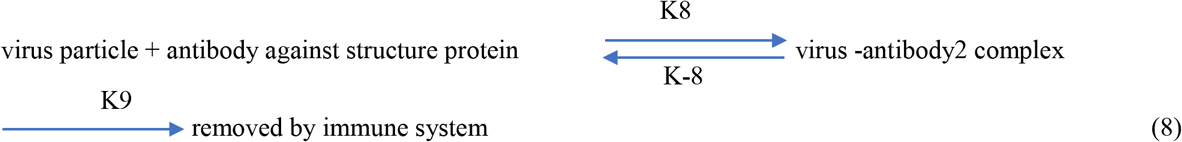

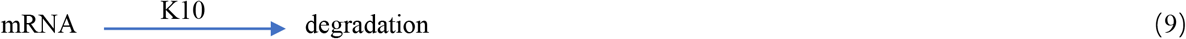

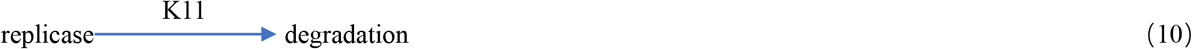

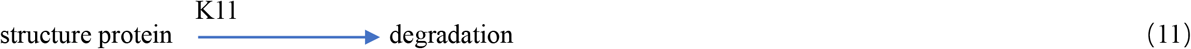

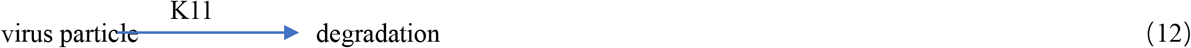

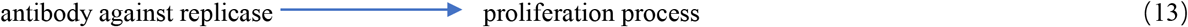

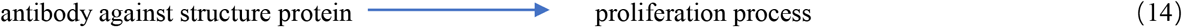

Reaction (1) is the replication process of virus, reaction (2) and (3) are the translation process of virus mRNA, reaction (4) is the packaging process of virus, reaction (5)-(8) is the interaction process between antibody and virus, and reaction (9)-(12) is the degradation process of different components of the virus. Reaction (13) to (14) are the antibody regeneration processes. We think that the evolution track of the virus is the process of maximizing the production of virus particles because only complete virus particles are infectious, and neither single mRNA nor protein is infectious. The ordinary differential equations based on those reactions are listed below:

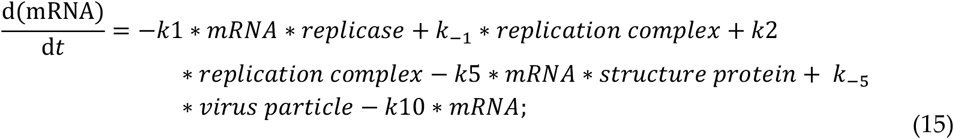

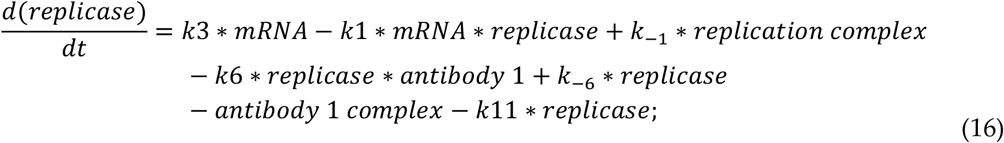

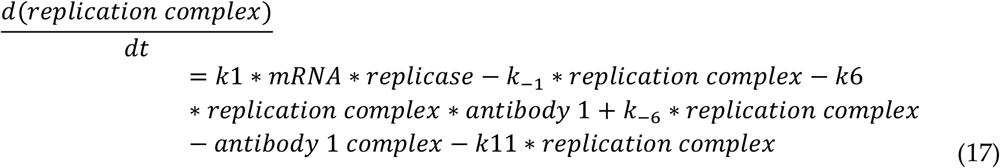

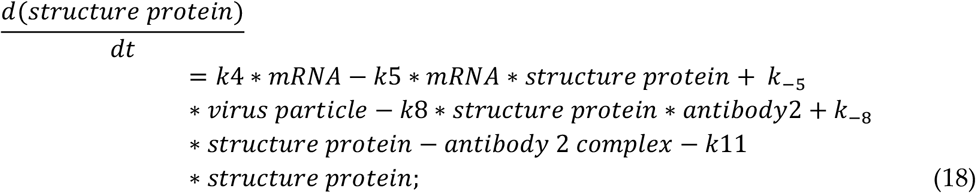

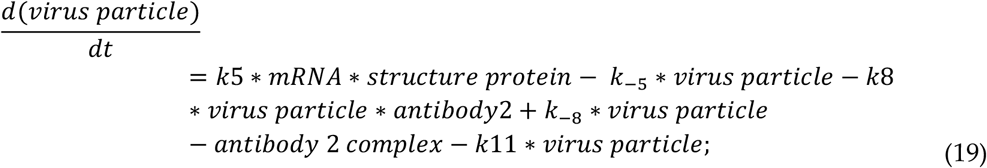

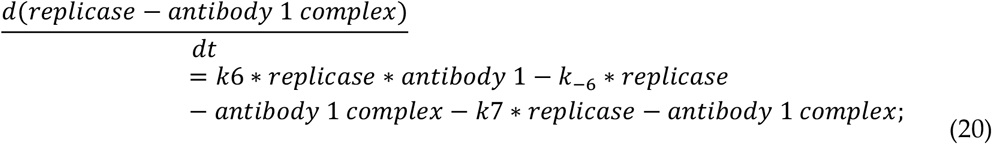

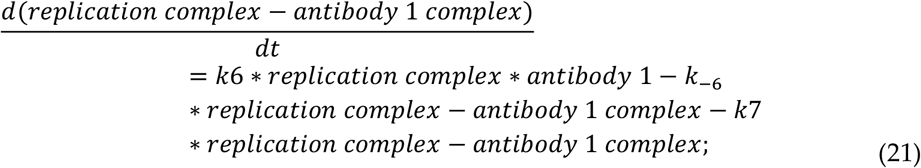

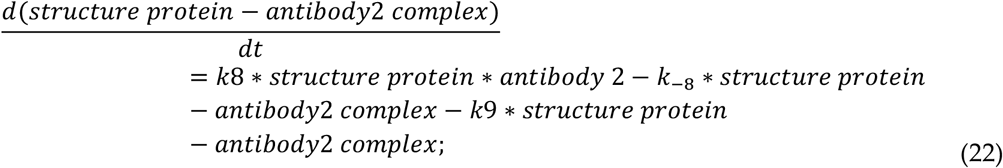

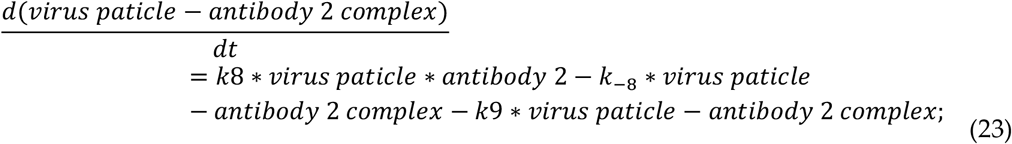

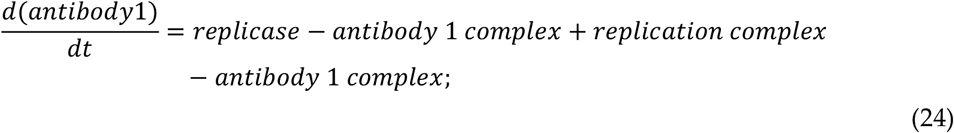

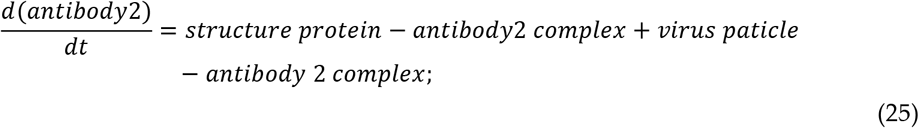

Equation (24) and (25) are used to describe the antibody proliferation process. The antibody dynamics model was firstly proposed in our previous research [21]. Here, for simplicity, the waning effect of antibody was not taken into consideration. The proliferation of the antibody was linearly correlated to its binding complex since this binding complex will further stimulate the regeneration of specific antibodies. We treat the immune clearance of antibody binding complex is very fast and efficient. Therefore, k7 and k9 was assigned to 1 in the following simulation process. Meanwhile, we proposed that the degradation rate was significantly different between mRNA and proteins. We assigned a larger value to k10 while keeping k11 as a small number.

### 2.2: Introduction of our small-scale model (model 2)

To better illustrate how mutation can dramatically influence its toxicological characteristics, a kinetic model that simulating the virus packing process is constructed. The agglutination nucleus is treated as the mRNA. In general, the virus packing process can be represented as multiple capsid proteins assembling around their core mRNA. This would eventually lead to the formation of a complete virus particle. The virus assembly process is a highly complicated process which might include the induction of signaling molecules, the formation of assembling nucleus and the final addition of membrane proteins or spike proteins outside. Here, we only model the core part, which is the assembling of capsids and their binding with mRNA. The processes are represented as the following reaction:

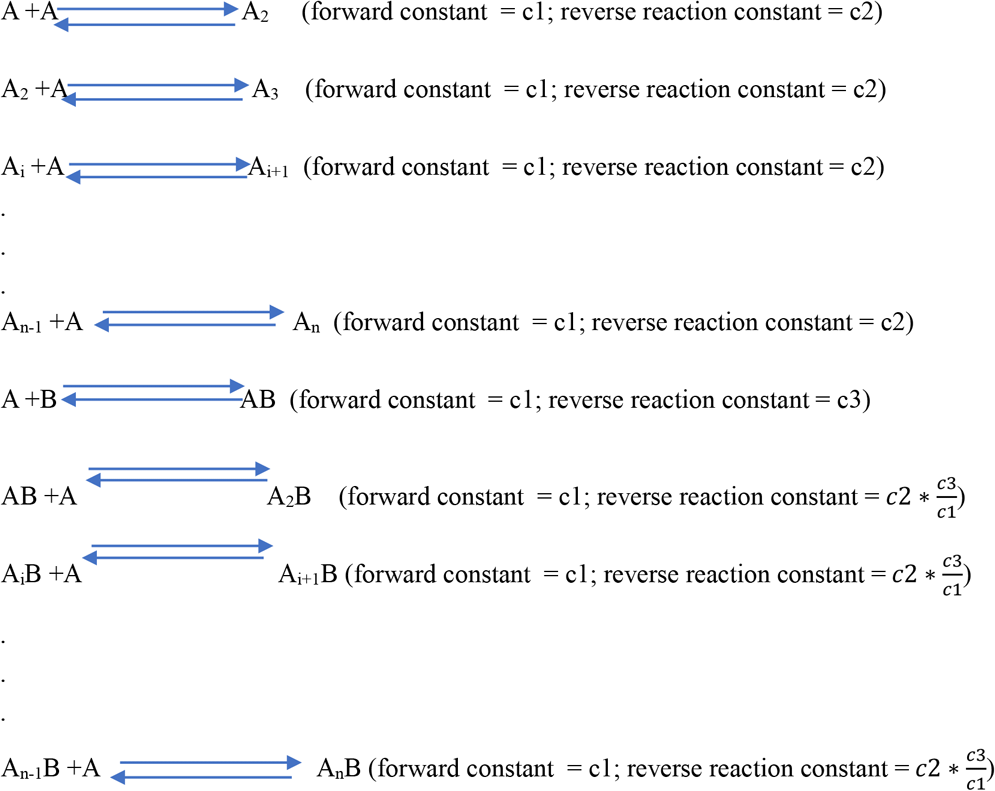

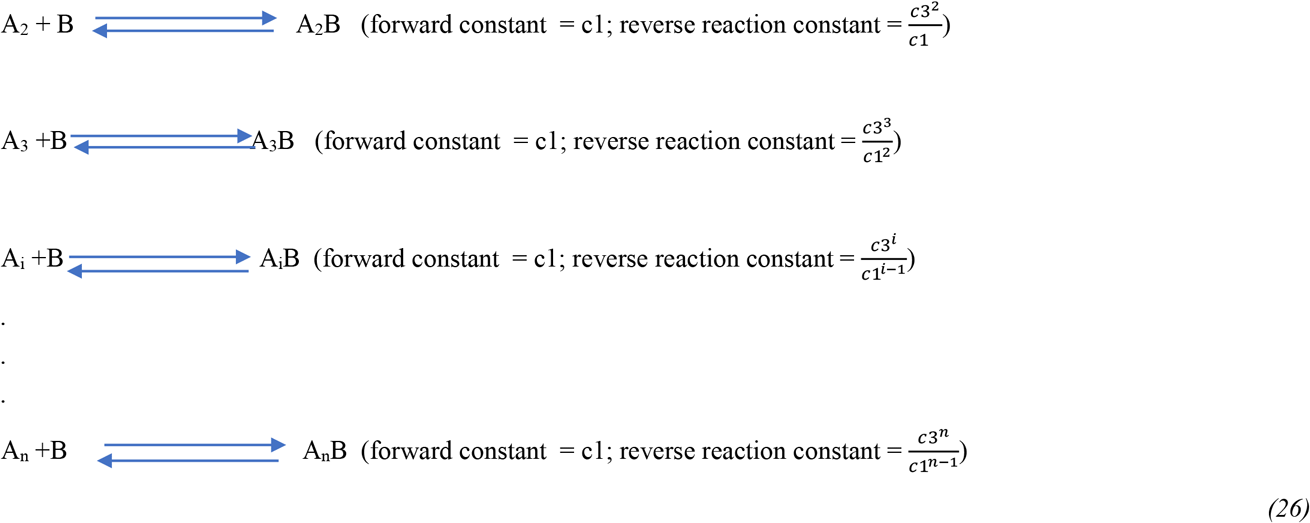

The ordinary differential equations based on those reactions are defined as below:

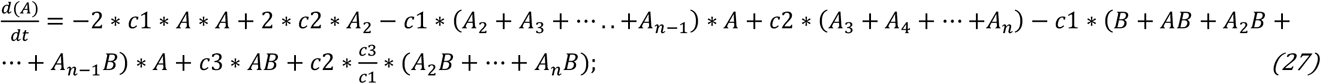

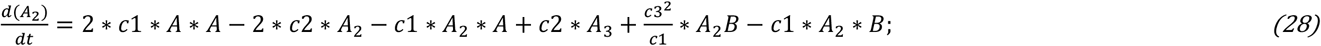

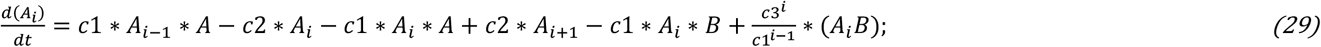

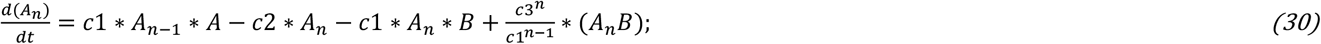

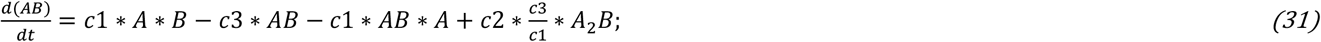

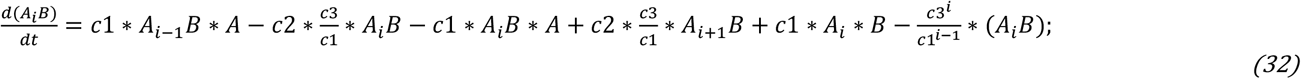

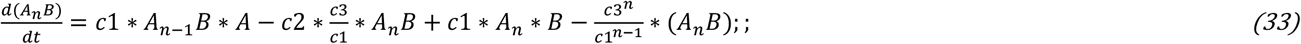

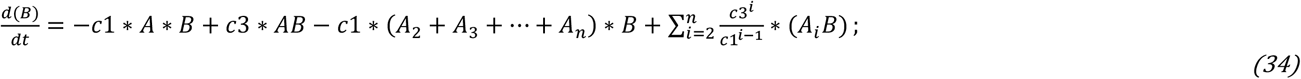

Here A represents the capsid or structure protein. An is the polymer that contains n structure proteins. B represents mRNA. *A*_*n*_*B* represents the final packed virus particle which has n structure proteins and one mRNA inside. *A*_*i*_*B* represents the incomplete packing intermediates which has *i* structure proteins and one mRNA inside. All the equations above are numerically solved using MATLAB ode15s function [22]. In result section 3.1, we will explain in detail how we determine the reaction coefficient of each reaction in the process (26).

## 3 Results

### 3.1: A general description and explanation of our model

In general, two mathematical models are presented in the methods part. The first one is the coarse-grained model which is used to predict the overall evolutionary trend of the RNA virus. The second one is the kinetic model that is used to simulate the virus assembly process. The second model is used to illustrate why tiny number of mutations would lead to a significant change both at virulence and transmissibility. The definition of virulence and transmissibility in our model is firstly provided before we give a detailed explanation of the first model. The virulence of the virus can be treated as the peak generation of heterogenous proteins, which is normally proportional to the overall immune response. The immune response can be mainly featured by the overall generation of antibodies which bind to those heterogenous proteins. Therefore, the final antibody generation quantity can be used as a benchmark to evaluate the virulence, especially toward same virus infection. However, the transmissibility of virus should be defined as the overall generated virus particles during one infection cycle. While mutations in spike proteins can significantly influence its transmission capacity, another important factor that influences virus transmissibility is the overall virus particles generated during one infection case. Experiments have indicated that the new emerging SARS-CoV-2 strain have more virus shedding compared with its previous strain [23-26]. The overall virus particles amount is different from the overall heterogeneous proteins. In the first model, the coefficient in reaction (1) is not feasible to be altered. Besides the binding coefficient between template mRNA and replicase, the main driving force of virus replication is that this replication process needs other energy units such as raw materials and ATP. Therefore, the mutational effects on reaction (1) are very weak. Similarly, reactions (2) to (3) are related to the translation efficiency of proteins, and the probability of significant changes caused by a few mutations points is also small. Reactions (5) to (14) are also weakly affected by mutations. Therefore, reaction (4) is the most sensitive process toward mutations. This will be fully illustrated in our second model. Mutations will lead to a slight increment in the binding energy between capsid protein and mRNA, they might also lead to a slight increase among capsid monomers. This slight change in binding kinetics will lead to a significant difference in the final formation of complete virus particle. The distinguished point of our theory is that we proposed that the enhancement of packaging ability will greatly weaken its replication ability, because we can see from reaction (1) that only the naked mRNA can make further replication, while the mRNA wrapped by capsid protein cannot make further replication and translation. Therefore, the improvement of virus packaging ability will inevitably lead to the decline of virus replication and translation ability. It is the increase in the number of virus particles that leads to the decline of its reproductive capacity, followed by the decline of the overall mRNA level and the content of all translation proteins, which also corresponds to a relatively low immune response (lower antibody production). We believe that the toxicity of virus depends on the total amount of heterologous substances produced, that is, the sum of mRNA, replicase and structural protein. This toxicity can also be measured by the highest level of antibody generated by stimulation. With the improvement of virus assembly ability, the number of virus particles will be significantly increased, and the infectivity will be significantly enhanced, but its toxicity will be significantly reduced. However, this assembly ability will not be enhanced indefinitely. Take two extreme cases for example. The first extreme case is that the virus packing capacity is 0, which means that it is impossible to package into virus particles. In this case, the number of virus particles is naturally very small, almost zero. In other extreme condition when the assembly ability is super strong, and all mRNA will be wrapped by its structural protein to form virus particles. In this case, because there is no naked mRNA, the virus cannot be further copied and translated, so the number of virus particles finally formed is also very small, almost zero. Therefore, there is an optimal binding force or binding coefficient k5 and k-5 between these two extreme cases, which can ensure the maximum virus particles produced. We think that this is the driving force behind the evolution of virus. The original COVID-19 strain has a weak packaging ability, so it replicates faster and is highly toxic. Meanwhile, its virus particles are in low quantity, thus has a relatively low transmissibility comparted to late strains. With the extension of time, the strain with strong packaging ability will gradually replace the strain with weak packaging ability, because the number of complete virus particles produced by it will increase, but this packaging ability will not increase indefinitely, and there is an optimal packaging ability to produce the largest number of virus particles. In result section 3.2, we will illustrate how the change in packing capacity will influence the virus evolution.

To better demonstrate why tiny binding energy change between capsid monomer and mRNA, together with the slight improvement in binding energy among those capsid monomers, would lead to a significant change in the kinetic coefficient in reaction (4) in our coarse-grained model, we incorporate a second model to simulate the complicated assembling process. Here are some assumptions in the second model: each capsid protein has a single binding surface toward mRNA or other capsid protein. Therefore, the binding energy between A_n_ and A is same to the binding energy between A and A. According to the thermodynamics equation ln Kd =-ΔG/(RT) [27]. The dissociation constant c2/c1 will be same for all the reaction listed below:

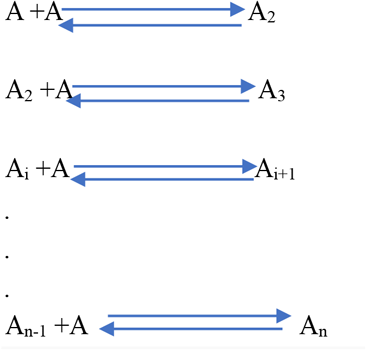

Since we assume the forward reaction kinetic constant c1 remains same for all reactions, all the reverse reaction kinetic constants are defined as c2.

For the reaction 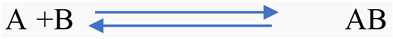, we use the same forward reaction kinetic constant c1. However, since the binding energy between capsid and mRNA is different from two capsid themselves, this reaction has a different dissociation constant. Therefore, a different reverse reaction kinetic constant c3 is used.

For the reaction 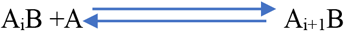, the binding energy can be classified into two parts: the binding between capsid and mRNA; and the binding among capsid themselves. Therefore, the dissociation constant of this reaction should be c2/c1*c3*c1. Since we treat the forward reaction constant as c1, the reverse reaction kinetic coefficient can be derived to be c2*c3/c1.

For the reaction 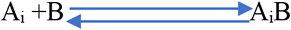, the binding energy is i times of the binding energy between A and B. Therefore, the dissociation constant of this reaction should be 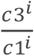. Since we treat the forward reaction kinetic as c1, the reverse reaction kinetic coefficient can be derived as 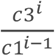. The assembly process of SARS-CoV-2 is normally contributed by hundreds of capsid proteins. Therefore, n should be a very large number. Here for simplicity, we use n = 5, 10 and 20 to perform the simulation in section 3.3. We illustrate the bigger the n be, the more significant the scale effect would be. A tiny alternation of binding energy among monomers could lead to a dramatic change in the final particle formed.

### 3.2 The coarse-grained model can predict the overall evolutionary direction of SARS-CoV-2

#### 3.2.1 A possible mechanism underlying the trade-off relationship between virulence and transmissibility

Parameters we used are: x0(1) = 0; % virus_spike_antibody1_complex

x0(2) = 0; % virus_complex_antibody1_complex

x0(3) = 0; % enzyme_antibody2_complex

x0(4) = 1; % initial virus particle (=mRNA + structure protein) level

x0(5) = 10; % antibody_1 which bind spike protein

x0(6) = 10; % antibody_2 which bind to enzyme

x0(7) = 1000; % initial mRNA level

x0(8) = 1000; % initial structure protein level x0(9) = 0; % initial enzyme level

x0(10) = 0; % initial enzyme_mRNA complex level

x0(11) = 0;% initial antibody2_enzyme_mRNA complex level

k1 = 1e-5; k-1 = 1e-5; k2 = 0.15; k3 = 0.5; k4 = 0.5; k5 = 1e-3; k-5 = 0.9; k6 = 5e-6; k-6 = 1e-14; k7 = 1; k8 = 5e-6; k-8 = 1e-14; k9 = 1; k10 = 0.1; k11 = 0.002.

We simulate the virus particle dynamics together with antibody dynamics in figure 1. As shown in figure 1, when the mutation increases the viral assembly capacity, the reverse reaction constant k-5 will decrease. The decrease in k-5 will lead to a smaller quantity of antibodies, which corresponds to a milder virulence. The overall particle number can be calculated as the cumulative virus number over time. Although the peak virus particle loading amount of the dashed yellow line is smaller than that of the dashed blue line, its cumulative number over time is larger than that of the dashed blue line. Therefore, mutations that promote the virus packing process will display an evolutionary advantage over its original strain. However, this trend is not continuous. As shown in the dashed green line, when the virus packing capacity surpasses certain threshold, a further decrease in k-5 will lead to a smaller cumulative number. The logic behind the trade-off between virulence and transmissibility in our model is that we assume the virus would optimize its shedding particle scale. Any improvement in the packing capacity will form an obstacle to its RNA replication and translation, thus decrease the overall heterogenous proteins. This will display a reduction in virulence in clinical performance. Before reaching the optimal point of maximizing the overall particle number, the improvement in packing capacity will lead to the generation of a more completed packed virus. This will display as a strong enhancement in transmissibility with a larger reproduction R_0_ value. After surpassing this optimal point, further mutations lead to faster packing will lose their evolution advantages since fewer virus particles would be generated. Therefore, the declination of virulence is not perdurable. The virulence of specific virus might engage significant declination at the early stage of the epidemic but would remain at a certain range after it reached the delicate balanced point. We proposed that this might be the underlying mechanism behind the intricate trade-off relationship between virulence and transmissibility.

#### 3.2.2 An optimal point or the balance point that maximizes the transmissibility can be theoretically calculated based on our model

If we fixed the rest parameters as described in section 3.2.1, we could get an optimal solution of k5 or k-5 in reaction (4). We could also calculate the overall virus shedding quantity under different combinations of k5 and k-5.

k5 can be calculated as 1e-6*1.05^(i), i is the value displayed in x-axis. k-5 can be calculated as 1*0.95^(j), j is the value displayed in y-axis.

**Figure 1:**
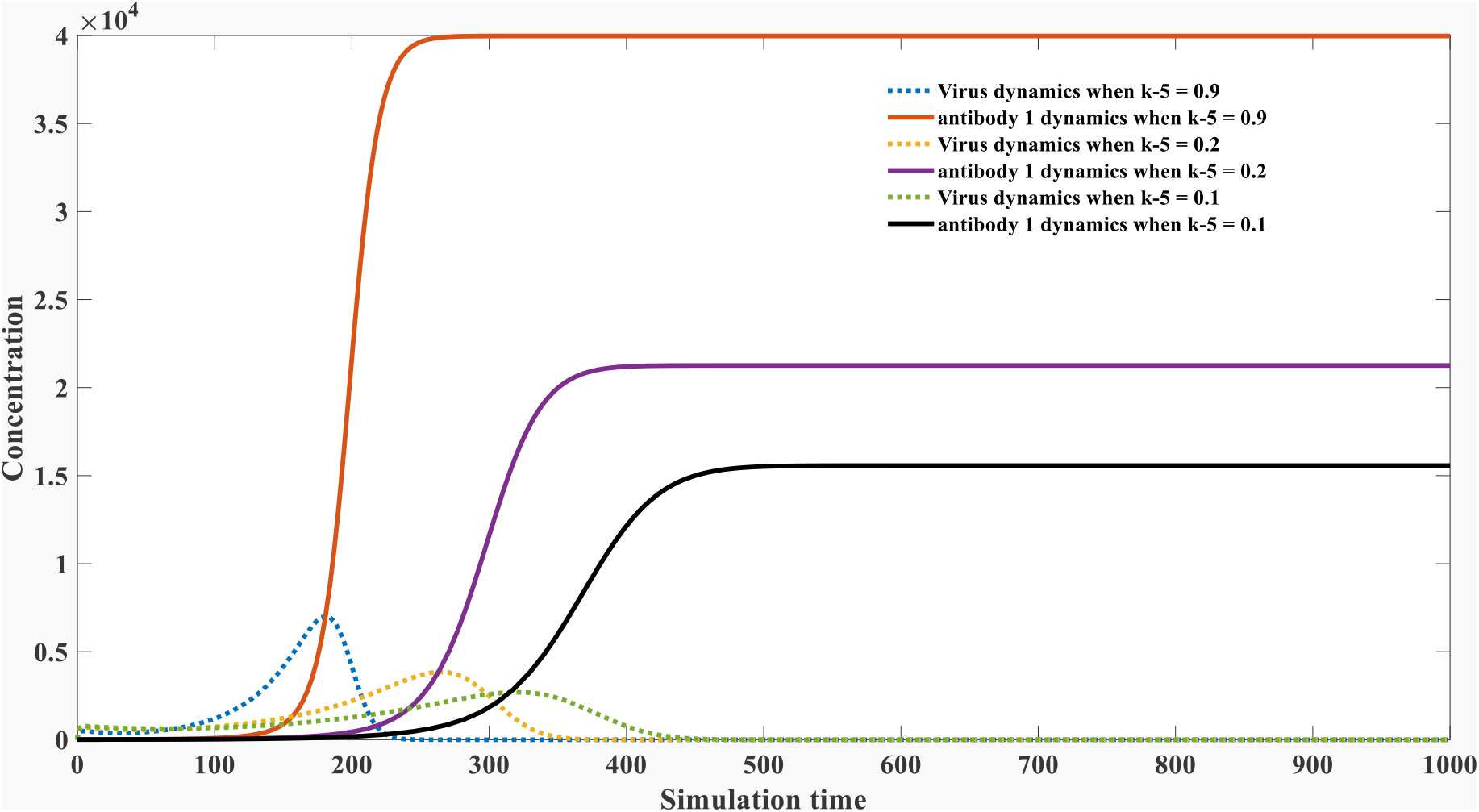
an illustration of the trade-off relationship between virulence and transmissibility.

It can be seen from figure 2, at the same k5 value, the overall virus particles generated are very sensitive toward the dissociation constant k-5. Mutations that promote viral assembly will lead to a significant increase in the overall virus formation quantity. When j increases in y-axis, k-5 will decrease correspondingly, the overall virus quantity will engage a significant growth before it reaches an optimal point. The forward reaction constant k5 also influences the overall virus particles number, but not as significant as k-5. The binding energy can be transformed into the equilibrium constant kd, which is equal to k-5/k5. We could obtain an optimal combination of k5 and k-5 using a global searching approach. We could also calculate the optimal kd value which is directly correlated with binding energy.

**Figure 2:**
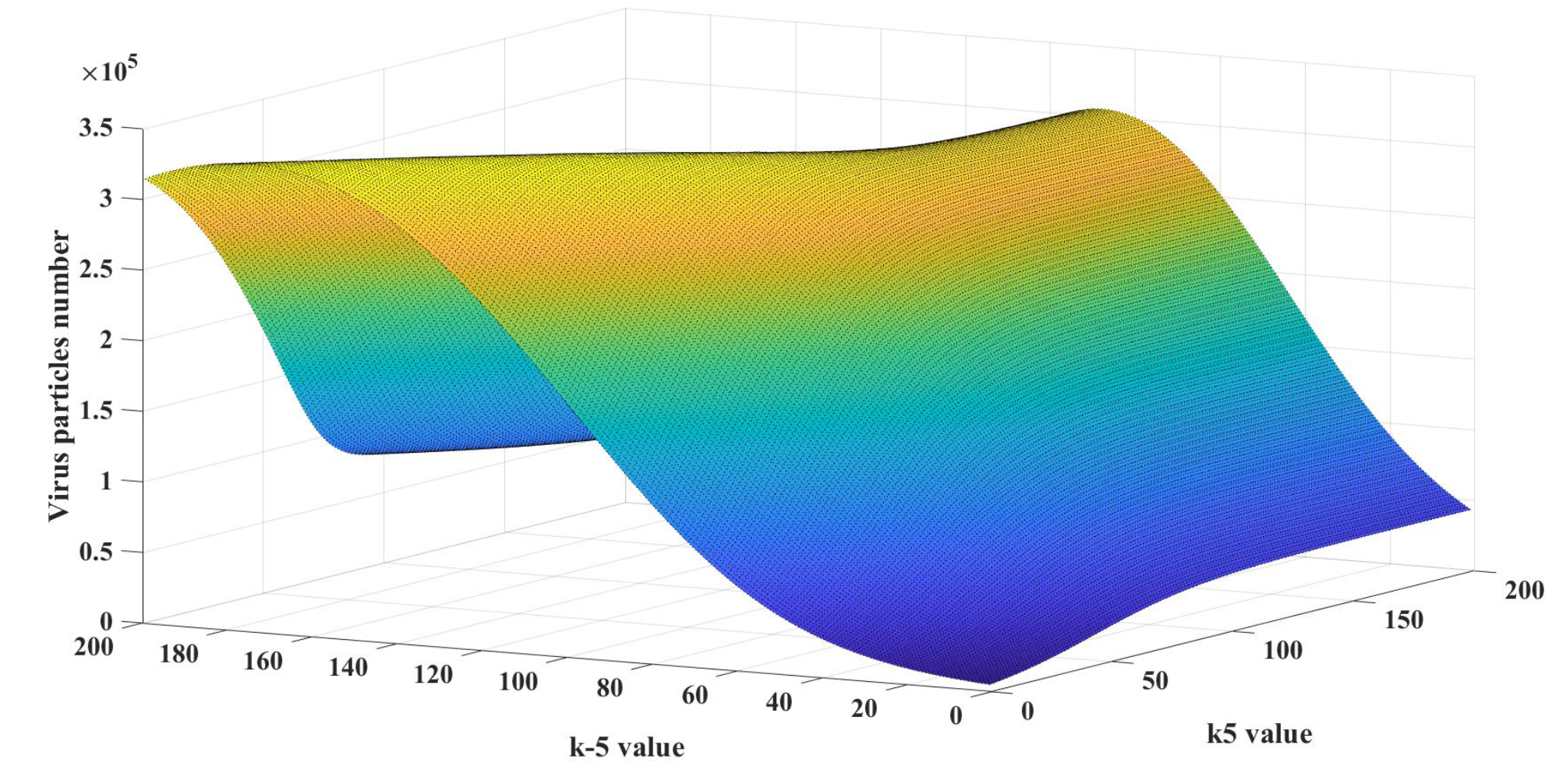
overall virus shedding amount at different k5 and k-5 value.

#### 3.2.3 Initial specific antibody level will influence this balance point

In section 3.2.1 and 3.2.2, we mathematically demonstrated that there was a balance point in which maximal virus particles could be generated. Before this balance point, mutations promoted the viral assembly will be evolutionarily selected and their virulence would display declination. In this section and section 3.2.4, we study the factors that influence this balance point. The initial antibody level might have an important influence on its virulence evolution.

We can see from figure 3a, the optimal value of k-5 will increase under a higher initial antibody level. This means massive vaccinations and infections will promote the virus to evolve into a more toxic strain. The viral assembly capacity will decrease to generate more complete virus particles in this case. However, it does not mean it is wrong to elevate our antibody level against SARS-CoV-2. Although the virus might evolve into a more toxic strain, its clinical features are determined by two parts: its own virulence and the immune response capacity from the host body. The elevation of the initial antibody level will surpass the virulence enhancement; thus, the overall virus particles and the general heterogeneous proteins will decline, as shown in figure 3b.

**Figure 3a:**
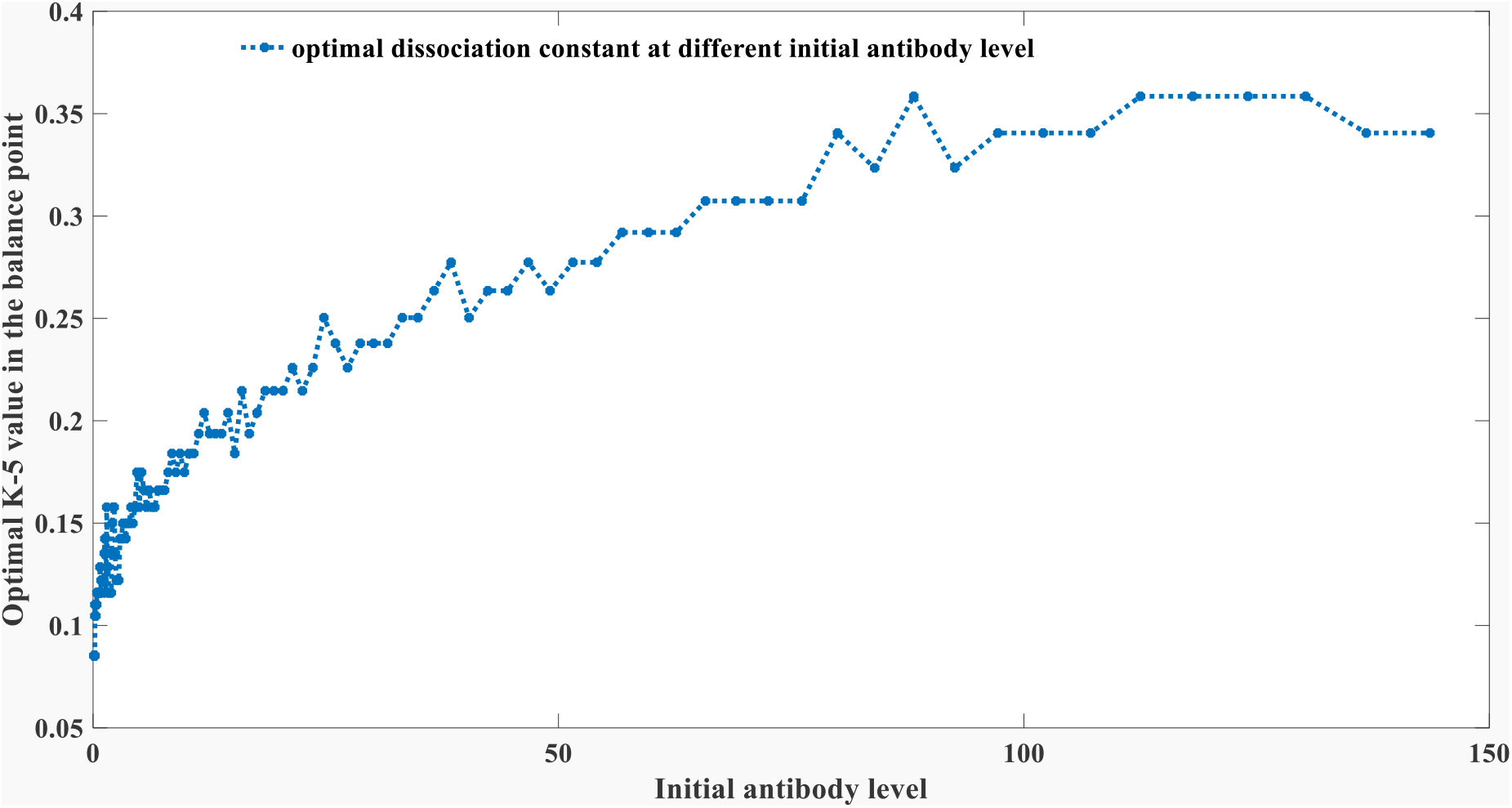
the optimal value of k-5 is different under different initial antibody levels

**Figure 3b:**
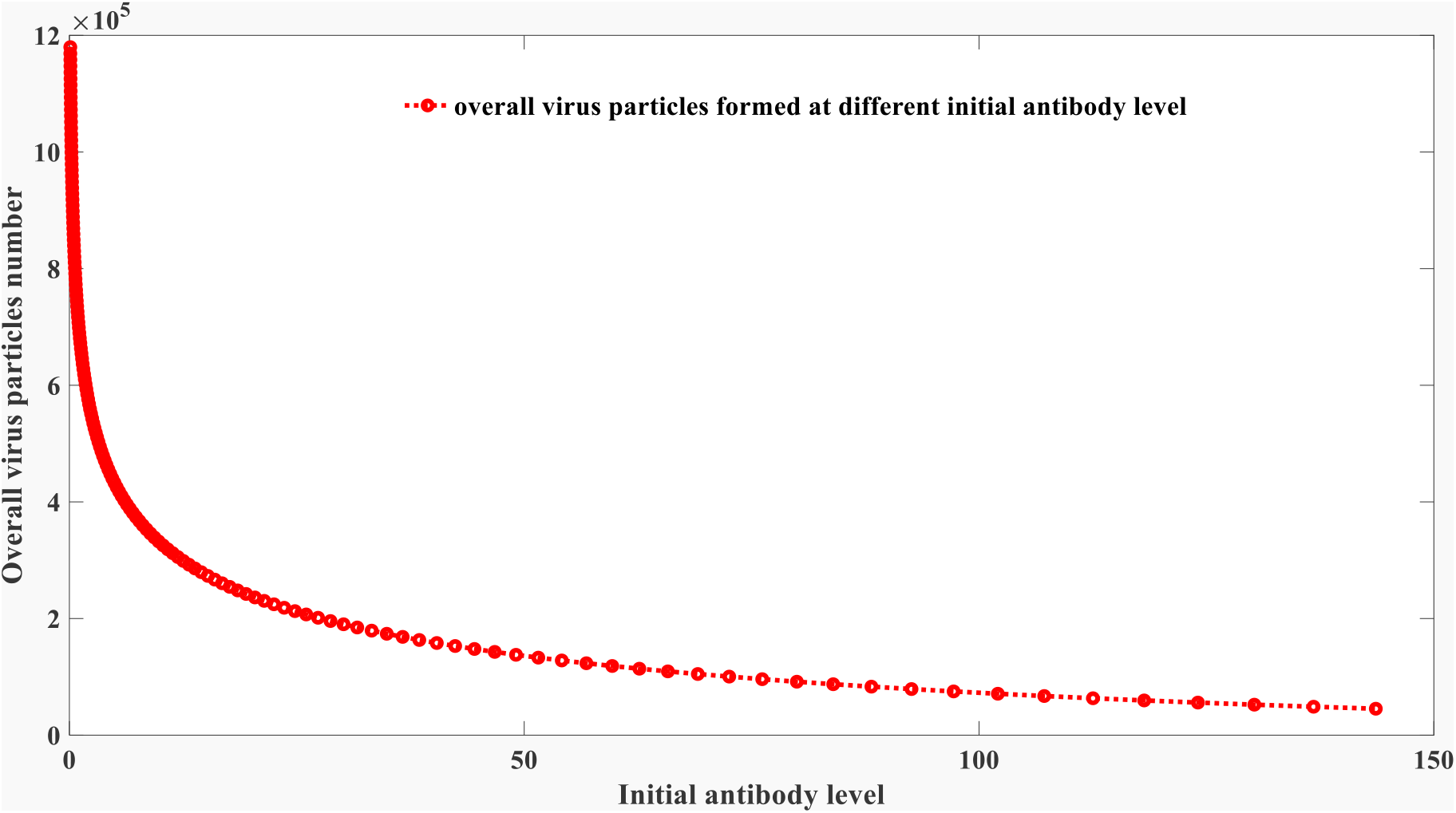
the overall virus particles number at different initial antibody levels

However, this might form a risk for other regions where few vaccinations or infections occurred. The virus evolved in a high vaccination area might lead to a much higher mortality rate in a low vaccination area because its virulence is evolutionary enhanced.

#### 3.2.4 Antibody binding features in different species will cause different optimal points in different species

Besides the antibody levels studies in the previous section, the antibody attribute would also significantly impact the final optimal equilibrium point. We supposed the binding capacity has a positive relationship with k-5 value. In species with fast binding kinetics, their k6 and k8 value are more prominent than human beings. In this case, the evolution process will select the strain with weak assembly capacity with an enormous k-5 value. The assembly capacity tends to be strongly truncated in species with suitable antibody binding kinetics such as bats. We gradually recognized the unique aspects of antiviral immune response in bats [28]. Besides the differences in the specific immune response, since bats are composed of small amounts of cells, this small-size mammal might have quicker binding kinetics against infection. Their antibodies are more feasible to get access to heterogeneous substances. As proposed in our previous publication, the forward binding constant is much more important than the overall binding affinity against antigen [21]. Therefore, bats should have a bigger k6 and k8 value than human beings. We also evaluate the influence of k-6 and k-8 on the optimal packing point, and we found the consequences of those reverse dissociation constants are very tiny. The modeling results are different from our expectations. As shown in figure 4a, the decrease of k6 could disfavor the virus packing process, enhancing its dissociation rate. The optimal dissociation constant k-5 will increase first, as shown in the green cycle marked as region 1. After it surpasses a threshold, it would display an opposite trend, as shown in the red area marked as region 2. We predict the cross-species transmission of coronavirus such as SARS, MERS, and SARS-COV-2 in region 2. The transmission from strong immunity species to weak immunity species will lead to a fatal pandemic in the new species. This can be shown in figure 4b, as the overall virus particles together with the peak antibody level significantly increase as the declination of the k6 value. However, as we observed in the epidemic development of SARS-CoV-2, its virulence was gradually decreasing. This demonstrated that the inter-species transmission of SARS-CoV-2 happened in the second region. The coronavirus with weak packing capacity is fatal to the species with slower binding kinetics. However, the immune features in the human population will promote the virus to evolve into tightly packed mutants. This will lessen its virulence at the same time. The attribute of the human immune system would gradually tame the coronavirus into a less virulent strain. Figure 4 can also explain why small mammals such as rats or bats are the original host of many viruses. They usually have stronger immunity with a fast antibody response kinetics, and their k6 value is located within the first region. The virus infection was not fatal to those animals since they have excellent immune kinetics, as shown in figure 4b. However, the virus’s packing capacity will be evolutionarily selected into a weak one. As shown in figure 4a, a larger optimal k-5 value will evolve in region 1. This fragile assembly virus is more virulent. It has a faster replication and translation speed since it has more naked mRNA during infection. This forms a significant threat to the relatively weak immune community such as our human beings. Therefore, for SARS-CoV-2, it might be possible that a more virulent strain is emerging in human society, but it is doubtful to evolve into a major strain under the evolution pressure. Meanwhile, we should concern about the inter-species transmission of new coronavirus from other species. This cross-species transmission might lead to a new epidemic outbreak as happened in SARS, MERS and COVID-19.

**Figure 4a:**
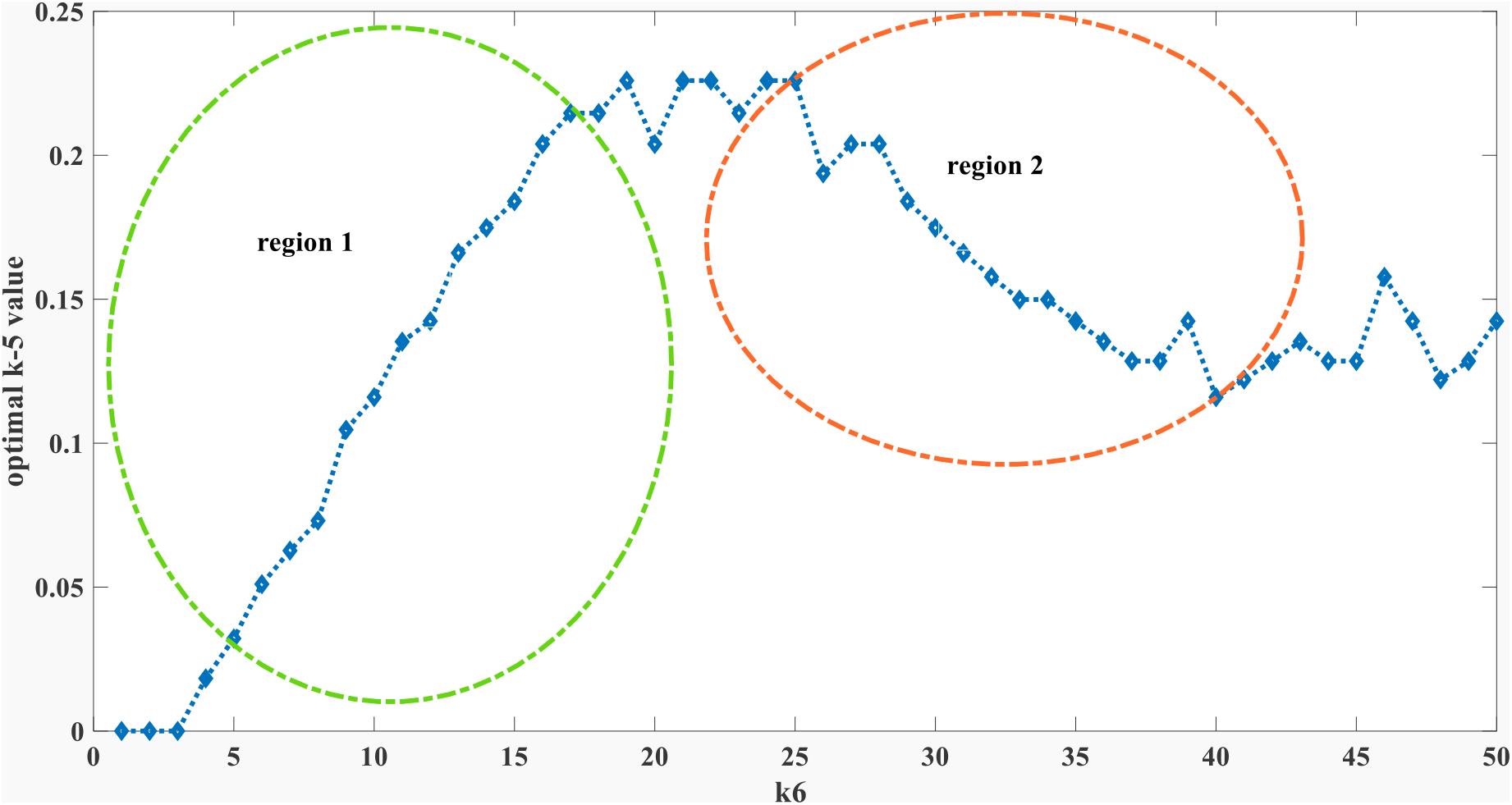
Relationship between the optimal packing capacity k-5 and antibody kinetic attribute k6. k6 is calculated as k6 = 1e-5*0.95^(i-1), where *i* is the number in y axis.

**Figure 4b:**
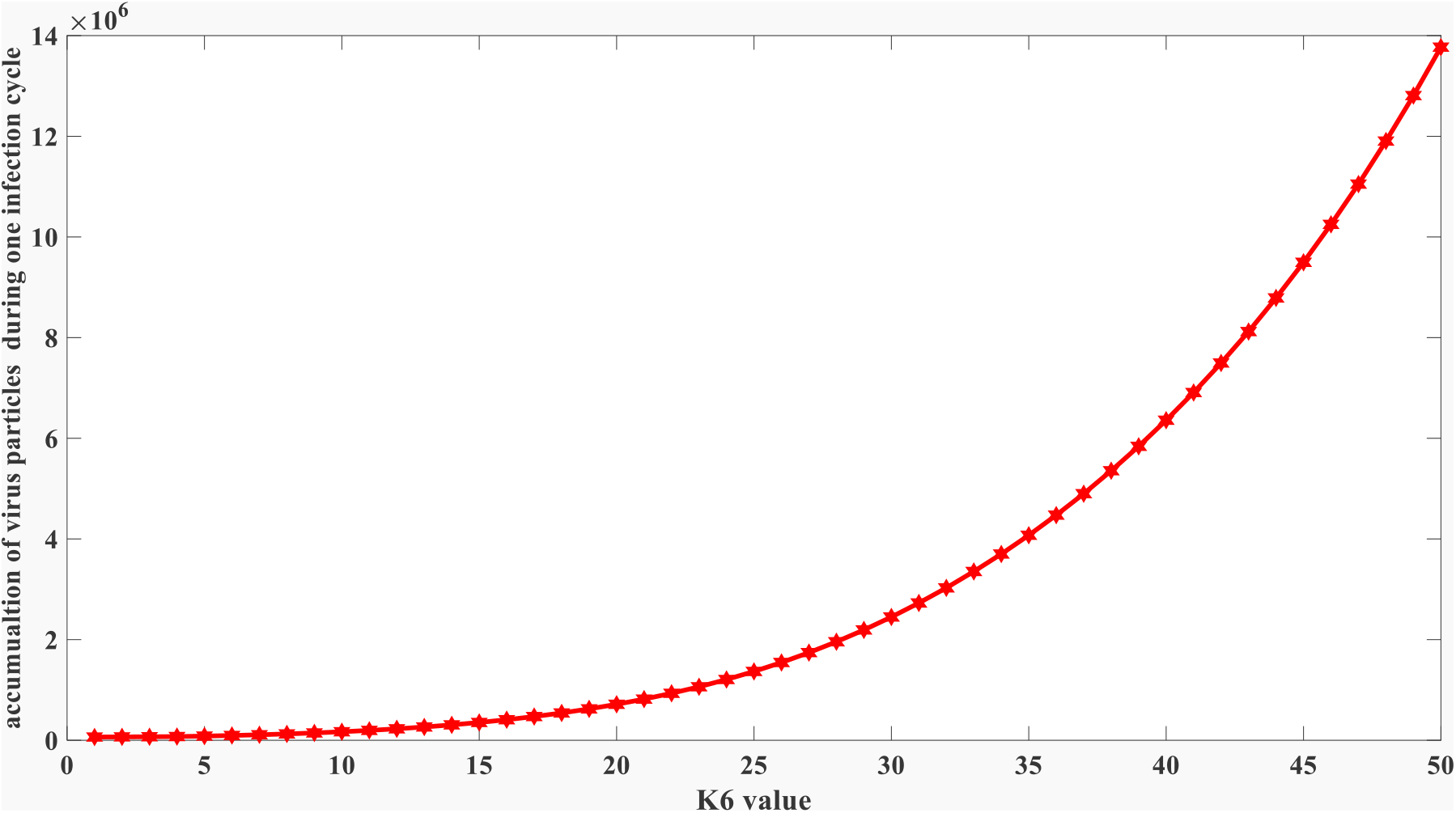
Overall virus accumulation quantity at different k6 values. k6 is calculated as k6 = 1e-5*0.95^(i-1), where *i* is the number in y axis.

### 3.3 The small-scale model can explain why a few mutations would lead to a significant change of its clinical features

The coarse-grained model is just a simplified representation of an actual situation. Although it reflects the subtle relationship between virulence and transmissibility, people might argue that it is not reasonable to have such dramatic change in k-5 value based on thermodynamic principles. For instance, according to the equation ln kd = -ΔG/(RT), the ΔG only engages minor changes due to mutation. Therefore, kd value and k-5 value cannot be altered in large magnitude. We have to reiterate that the coarse-grained model is not physically linked to the kinetics behaviors. We developed a small-scale model that can be physically connected to the binding energy perturbation due to mutation to simulate the viral assembly process better.

#### 3.3.1 our virus assembling model has a good match with experimental reports

The self-assembly of a set of viral proteins into a regularly shaped capsid involves a fine-tuned range of interactions between the capsid proteins to create this highly ordered, symmetrical, supramolecular structure. In a simplified view, assembly is a spontaneous process driven by weak protein-protein interactions on the order of several *kBT* [29]. These weak attractive interactions are mainly hydrophobic [30-31] and can overcome the entropic penalty of forming highly organized capsid structures. Coarse-grained models indicate that the growth phase proceeds as a downhill process, while completion is rate-limited due to steric effects [32-34]. The latter is even thermodynamically unfavorable when only looking at the loss in flexibility upon capsid closure [34]. Under assembly conditions, small intermediates are transient, and nuclei formation are rare events [35-36], making them challenging to characterize.

On the other hand, large intermediates are reasonably stable and easier to identify. However, a large number of protein subunits per capsid (for icosahedral capsids typically specific multitudes of 60, as described by the quasi-equivalence theory and the corresponding triangulation number [37])leads to limitations in the experimental resolution. It is hard for such large numbers of subunits to discriminate complete capsids from capsids missing a few subunits. Indeed, nucleation and completion are challenging processes to investigate, which explains why only a limited number of experimental studies have been reported on this topic. Another characteristic of virus self-assembly studies is that assembly/disassembly reactions are typically triggered under controlled conditions. The capability of CPs to assembly is frequently tuned by changing solvent conditions such as pH, salt concentration, or mild concentrations of denaturant agents.

The in vitro assembly and disassembly of one of the simplest virus particles known, the minute virus of mice (MVM), has been recently characterized using a combination of atomic force microscopy (AFM) and transmission electron microscopy [38]. MVM forms T = 1 icosahedral capsids of 25 nm made of only 60 CPs, which arrange as trimeric subunits. The whole range of particle sizes was found in this study [38], from complete capsids, via large incomplete capsids, to small capsid intermediates (Figure 5). Full capsids were analyzed separately according to their capability to be permeable to uranyl ions, as characterized by TEM experiments. The right graph in Figure 5 shows the assembly evolution of intermediates to complete capsids over time. In all cases, large and small intermediates are populated in low numbers and, hence, transient. This is indicative of a highly cooperative process.

**Figure 5:**
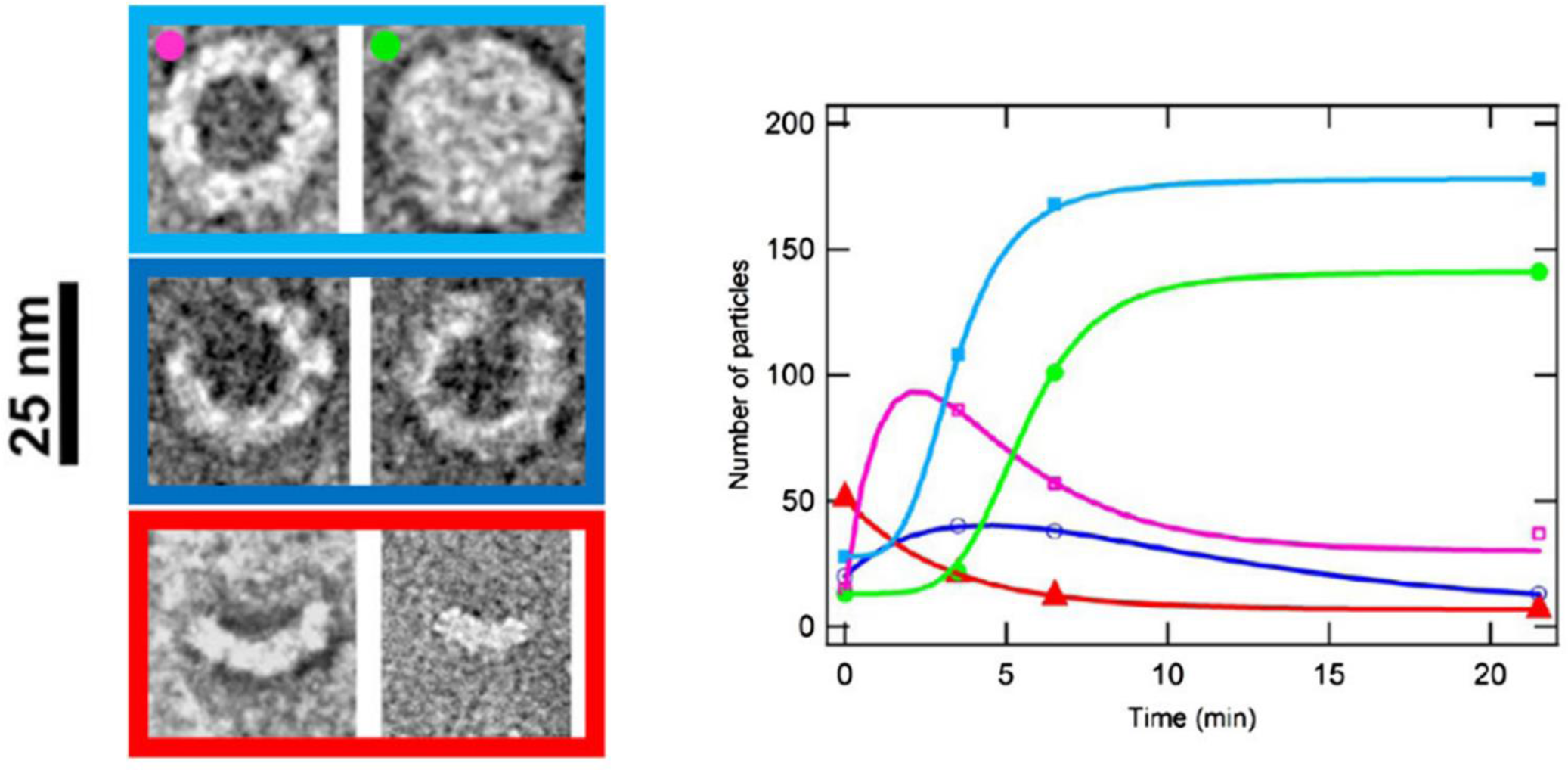
MVM particles imaged by TEM (left): light blue, Types I + II particles (complete capsids); green, Type I (complete capsids in basal state); magenta, Type II (complete rearranged capsids); blue, Type IIIA (large incomplete capsids); red, Type IIIB (smaller incomplete capsids). Progression of the total number of particles during assembly (right graph) over time. (Reprinted with permission from Medrano et al.).

Since the experiment in figure 5 only uses capsid proteins, we also removed mRNA in our modeling. As shown in figure 5, the complete packed particle will gradually increase through time. This trend can also be reflected in our model. The completely packed particle is represented as greed line in figure 6. It can be seen that the fully packed virus particle will also increase through time. Beside that, the poorly packed particles engaged similar declination as reflected by the experiment. The poorly packed particles encountered a transient increase and gradually declined after that, as shown in the purple line in figure 6. The well-packed particles have a similar trend compared to experimental reports. By comparing Figure 5 and figure 6, you can see our small-scale model has a good performance in reproducing the experimental results. This test adds more credibility to our kinetic model.

**Figure 6:**
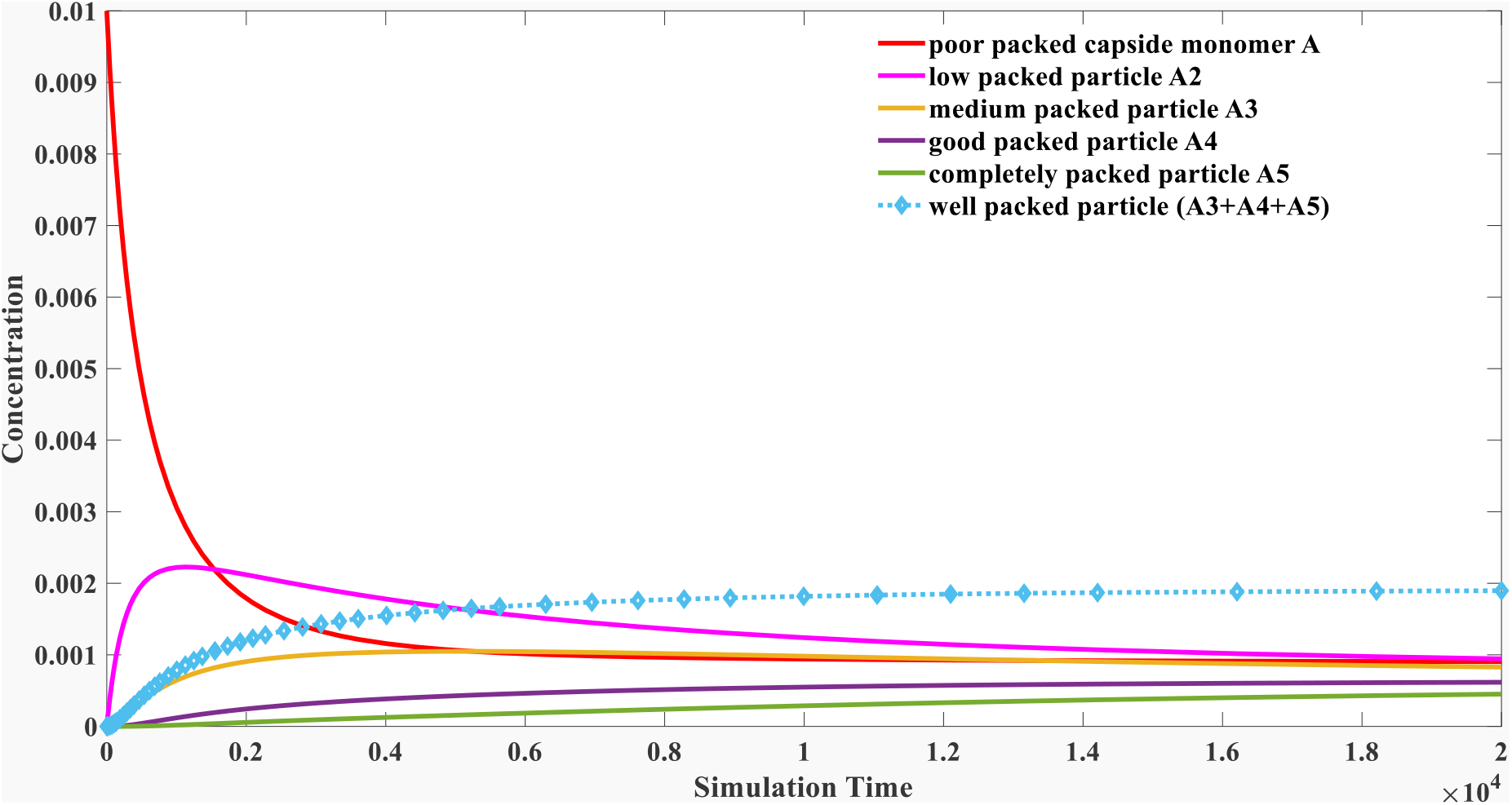
Simulation results of progression of the total number of particles during the assembly process over time in our small-scale model (n = 5 without mRNA).

#### 3.3.2 The small-scale model can explain why a few mutations would lead to a significant change in overall virus particle quantity

After the validation process, we try to explain why a few mutations would cause a significant alternation in the virus formed. We will illustrate how the number of capsid proteins in one particle influences the overall packing capacity. Three cases were compared in our study when we utilized n = 5,10,20 respectively. In all of those three cases, we think mutations would only lead to a minor enhancement toward the binding energy. Therefore, the dissociation constant c2 and c3 is only down-regulated by 10%. The results are shown in figure 7.

**Figure 7A:**
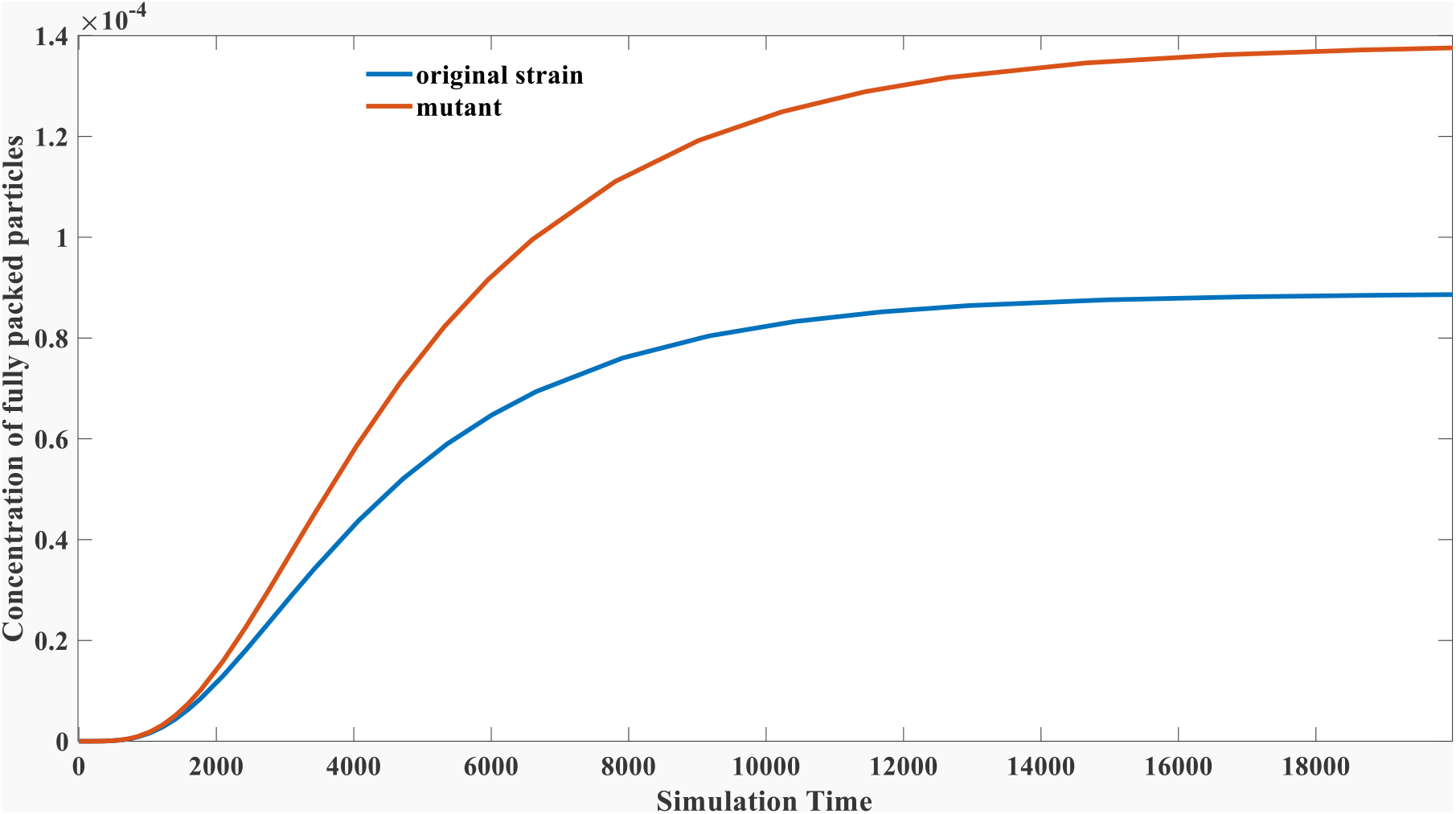
The impact of mutation on the total number of particles during the assembly process in our small-scale model (n = 5).

**Figure 7B:**
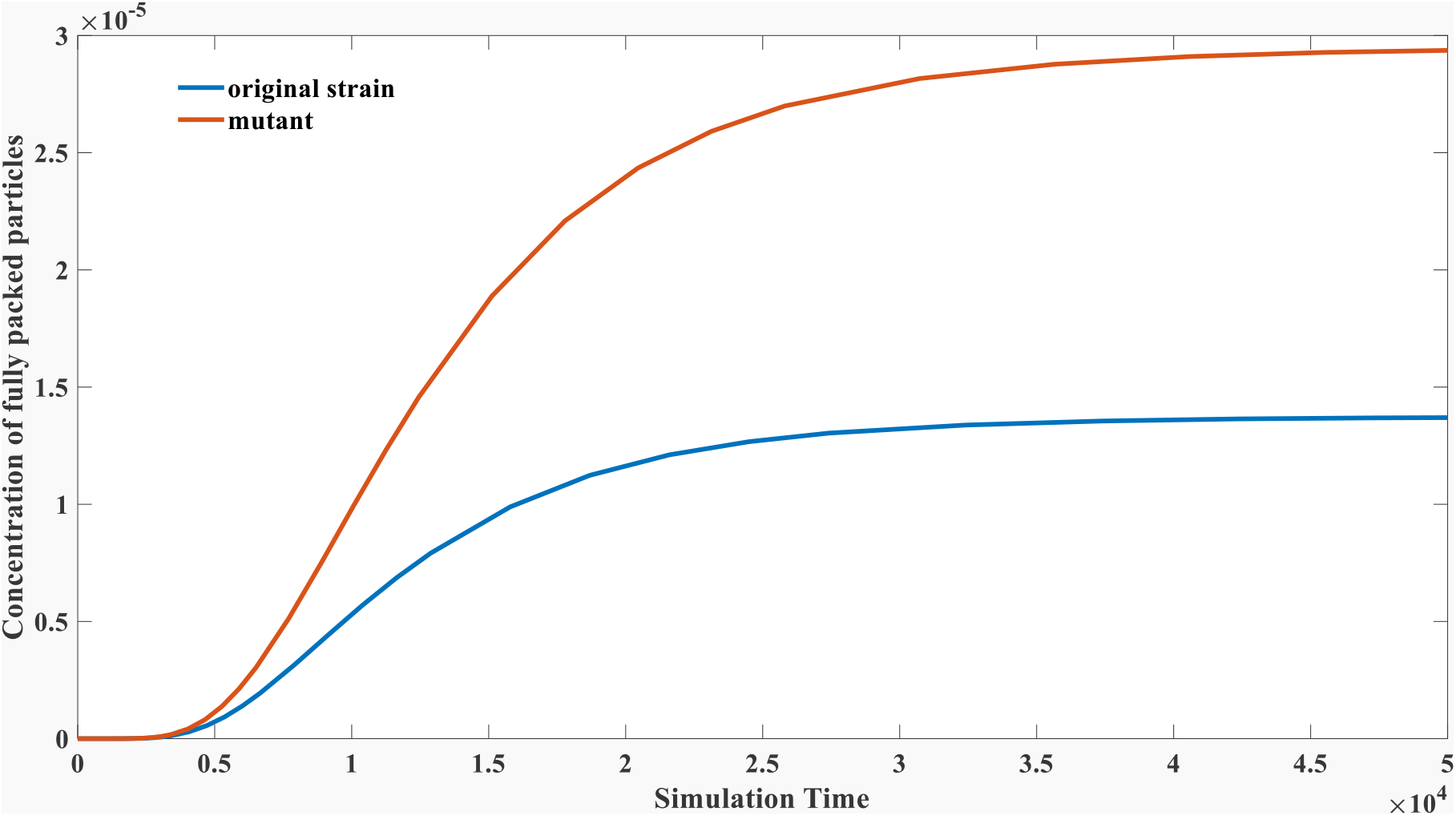
The impact of mutation on the total number of particles during the assembly process in our small-scale model (n = 10).

**Figure 7C:**
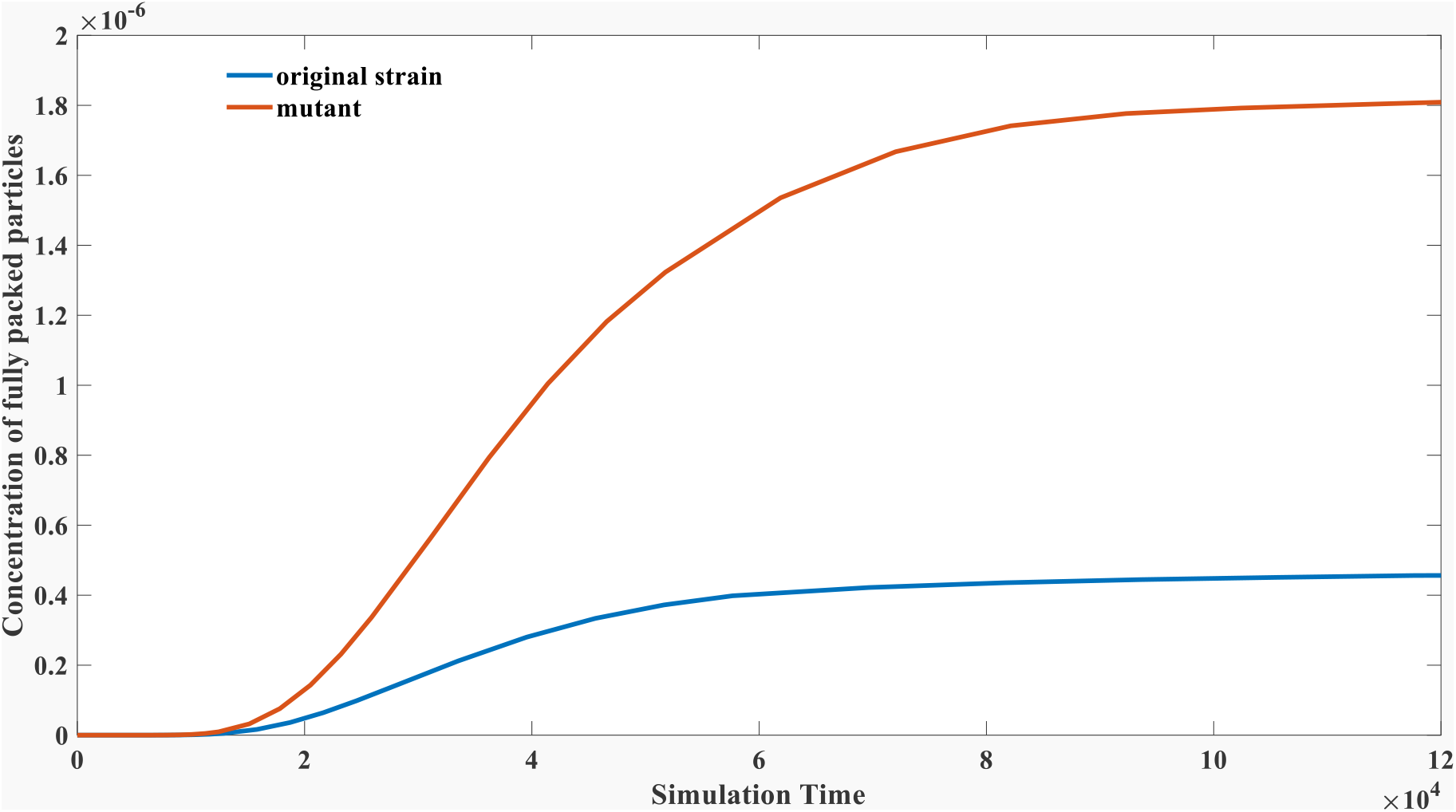
The impact of mutation on the total number of particles during the assembly process in our small-scale model (n = 20).

It can be seen from Figure7A that the mutation only causes 10% enhancement in binding affinity. The mutant strain displays a more robust particle formation capacity with a 50% increase than the original strain. When n increases, the magnification would be more significant as shown in figure 7b and figure 7c. For example, this enhancement was more than 100% when n = 10; and was 400% when n = 20. As we described above, the formation of SARS-CoV-2 particles typically recruits more than 400 capsid proteins [39-40]. Therefore, any little mutation effect in changing the binding behaviors of those capsid monomers or binding affinity between the capsid and mRNA would cause a significant alternation in the final quantity of virus particles. This can explain why some single mutation would cause a dramatic change in its clinical feature. This could explain why mutants such as omicron will shed viruses more than thousands of folds than the original strain.

#### 3.3.3 the small-scale model might explain why a few mutations would lead to significant declination in virulence

The declination in virulence might also be explained using the second model. It was noticed the newly emerged strain, such as omicron, displays weak virulence. It is hard to explain why small mutations would lead to a significant decline in virulence, especially in the early epidemic phase from March 2020 to May 2020. The overall death rate experienced a significant decrease, from 10% to 1% [41]. Although we cannot deny the possibility from other aspects such as UTR deletion proposed in our previous preprint publication [42], we explain that point mutation might also contribute to the rapid declination in virulence. The virulence in our second model can be reflected as the concentration of naked mRNA. We modeled the change of naked mRNA using the same parameter sets in figure7. The results are displayed in figure 8.

**Figure 8:**
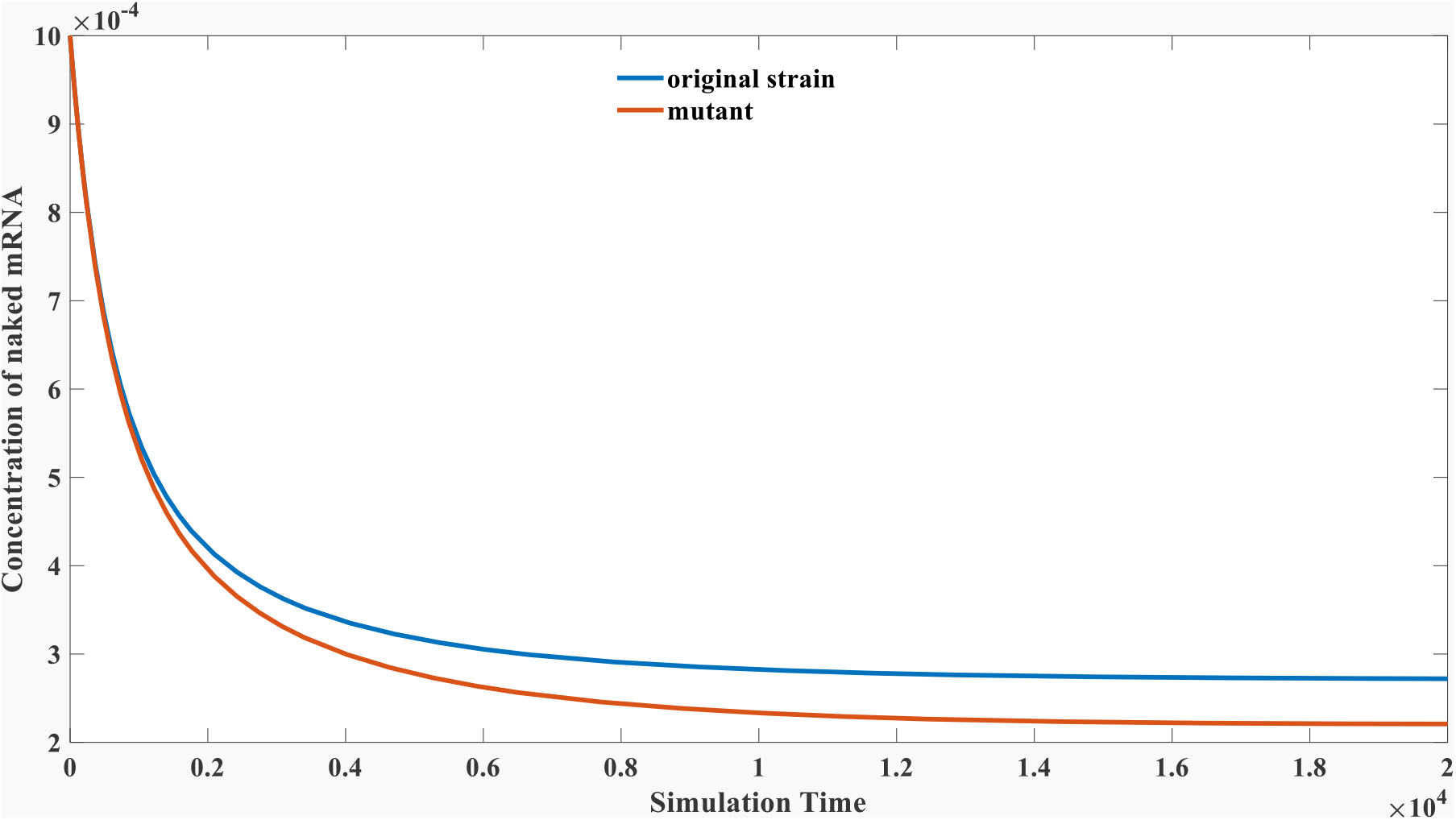
The impact of the mutation on the total number of naked mRNA during the assembly process in our small-scale model (n = 20).

It can be seen from figure 8 that the concentration of naked mRNA is decreased by 19% after mutation. Although this 19% is not significant, those naked mRNA will participate in further replication and translation. Therefore, a noticeable concentration difference in the equilibrium state might lead to a substantial difference after several rounds of replication.

## 4 Discussion

In the discussion part, we discuss the possible application of this theory, especially toward those sensitive questions listed in the introduction part.

### I: What is the exact relationship between virulence and transmissibility?

According to our coarse-grained model, there would be a balance point between virulence and transmissibility. The evolution will advance the formation of mutants that generate more virus particles. The key feature of our coarse-grained model is that we focus on the virus packing process on the overall virus formation. It is rarely noticed before that the strong assembly kinetic might inhibit the proliferation of new mRNA and proteins. The physical background of this model is that we think the mRNA is only active in the non-binding state. The binding of the capsid will hamper its replication and translation. Our simulation results indicate that the mutant with increased assembly capacity will be selected given an impoverished packing background. The virus shedding amount will engage significant enhancement, and its transmissibility will boom, as displayed in the case of SARS-CoV-2.

Meanwhile, as more mRNA are trapped around capsid protein, its replication capacity will decrease, displaying a weak virulence in the clinic. This trend can also occur in other RNA virus infections such as the 1918 flu virus, H1N1, myxoma, and the virus causing swine flu. The phenomenon described at the beginning of the introduction implied that almost all RNA virus epidemics engaged a virulence declination process, is not a historical coincidence.

### II: Do we need to worry about the emergence of a more deadly strain?

This question is critical since it greatly influences our prevention policy. While some scientists were concerned that a fatal mutant would eventually evolve [43], our model rejects this possibility. There might be some fatal mutants emerging, but the evolution pressure will limit the large-scale transmission of those toxic strains. Therefore, the overall evolutionary trend of SARS-CoV-2 is to deduce its virulence. Only by reducing their virulence can they increase their transmissibility. Consequently, we should not worry too much about the emergence of a more deadly strain since it is strongly impeded in evolution. However, as we pointed out in the result section 3.2.4, we are very likely to experience a new epidemic if another cross-species transmission occurs in the future. A novel virus instead of SARS-CoV-2 more likely causes this future epidemic.

### III: Will the virulence of SARS-CoV-2 finally fade away?

Although our model opposes the thought that a fatal strain would eventually emerge, we should not be too optimistic about the epidemic situation. According to our coarse-grained model, the virulence of SARS-CoV-2 might not be faded away. It might evolve into a less virulence strain as the evolution of 1918 flu. There will be a balance point as displayed in our model. Beyond this optimal point, a mutant with less virulence is unfavorable in evolution. The declination of virulence cannot be continuous. Unluckily, our model cannot accurately estimate this balance point. Nevertheless, it can be inferred that the optimal virulence at the balance point might be the same or even weaker than the omicron strain as clinical data revealed the emergence of new omicron sub-strains with more transmissibility less morality rate [44-45].

### IV: Why the cross-species transmission always leads to a fatal outbreak in the early epidemic stage?

While this study cannot entirely exclude the possibility of the artificial origin of SARS-CoV-2, it provides a rationality of the natural origin of this coronavirus. It could also help explain why cross-species transmission always results in a severe epidemic with high mortality in the outbreak stage. Section 3.2.4 demonstrated that the balance point is different in different species. For species with solid immunity and quick immune response, the dominant strain is inclined to have a worse packing capacity to maximize its overall particles formation. Therefore, the transmission from the bats’ community might cause a much high mortality rate in the human population. It might take a markable period to tame it into a mild strain under evolution pressure in the community of new species.

### V: Why a few mutations can significantly alter the clinical behaviors of SARS-CoV-2?

In section 3.3, it is mathematically demonstrated that a few mutations can lead to a significant change in the overall particle formation together with its virulence. While many researchers focus on the effect of mutation on spike protein which directly influences the virus entrance capacity, we suggested modifications on capsid protein or mRNA might play an essential role in determining its clinical features. Mutations that promote the binding among capsid proteins or the binding between capsid and mRNA would significantly enhance the final formation of complete particles. The polymeric aggregation process will amplify the tiny binding energy change into a tremendous difference in its assembly outcome. Therefore, it does not necessarily need many mutations to change its clinical severity dramatically. We should not evaluate its toxicity based on its homology and similarity to the original strain. For example, very high homology exists when comparing all strains of SARS-CoV-2. This does not indicate they have similar clinical performances. Some mutants, such as omicron strain, might be very mild, while the original template is fatal.

In the end, we have to admit there are many uncertainties in our model. This model is just a simple mathematical representation of a real scenario. The parameter values can be further improved and modified given sufficient experimental data. It needs further experimental or ecology validation eventually. Meanwhile, we did not intend to encourage people to reduce protection against SARS-CoV-2 when we acclaim the virulence of SARS-CoV-2 will engage a declination. This work only provides a hypothesis on the evolutionary relationship between virulence and transmissibility.

